## Supplemental Figures for "The landscape of metabolic pathway dependencies in cancer cell lines"

[†]Present address: Nautilus Biotechnology, San Carlos, CA

Running title:

Metabolic pathway dependencies in cancer

To whom correspondence should be addressed: Nicholas A. Graham, University of Southern California, Los Angeles, 3710 McClintock Ave., RTH 509, Los Angeles, CA 90089. Phone: 213-240-0449;

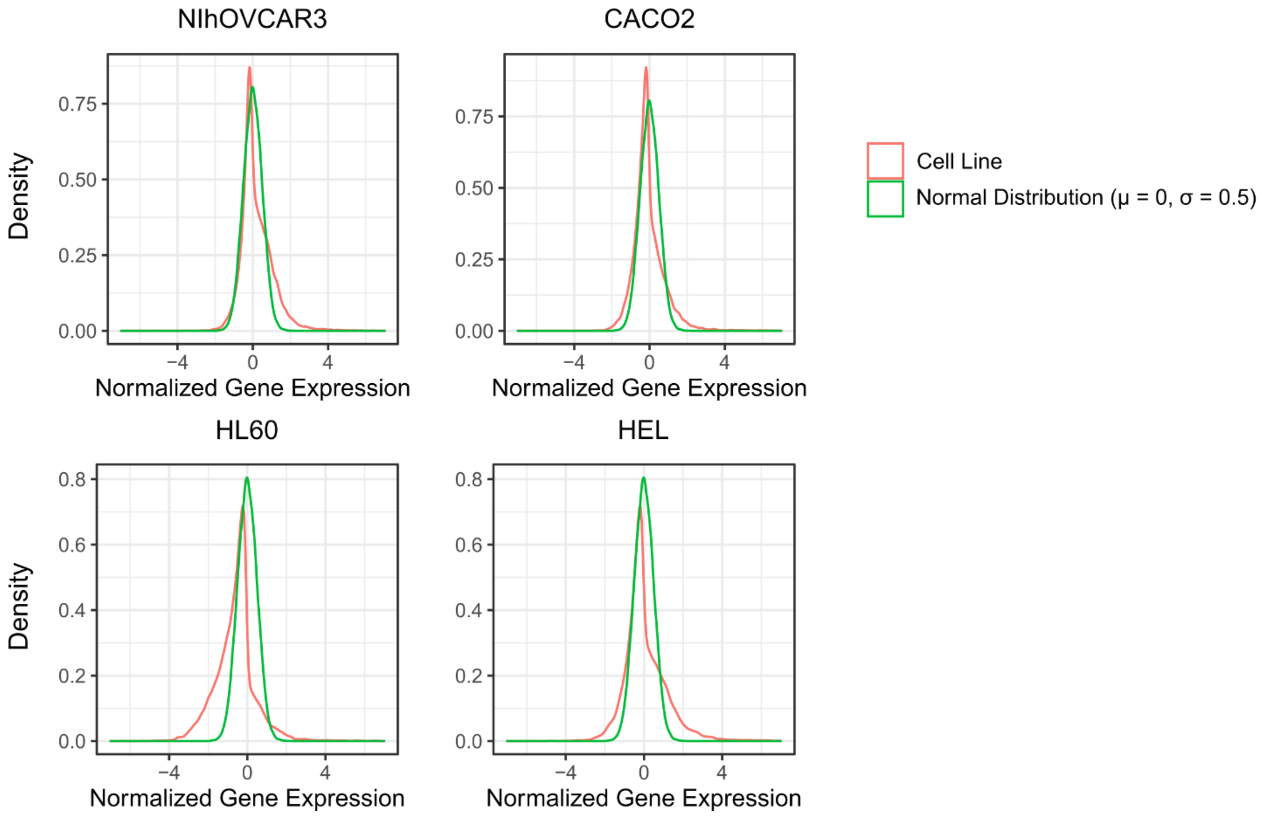


**Supporting Figure 1. A normal distribution with mean of 0 and standard deviation of 0.5 reflects gene expression profiles**. Gene expression data was taken from the Cancer Cell Line Encyclopedia (CCLE) and scaled and centered within culture type (adherent or suspension) and culture medium (DMEM or RPMI). Four cell lines were chosen at random and their gene expression profiles (red) were compared to a normal distribution with a mean of 0 and a standard deviation of 0.5 (green).

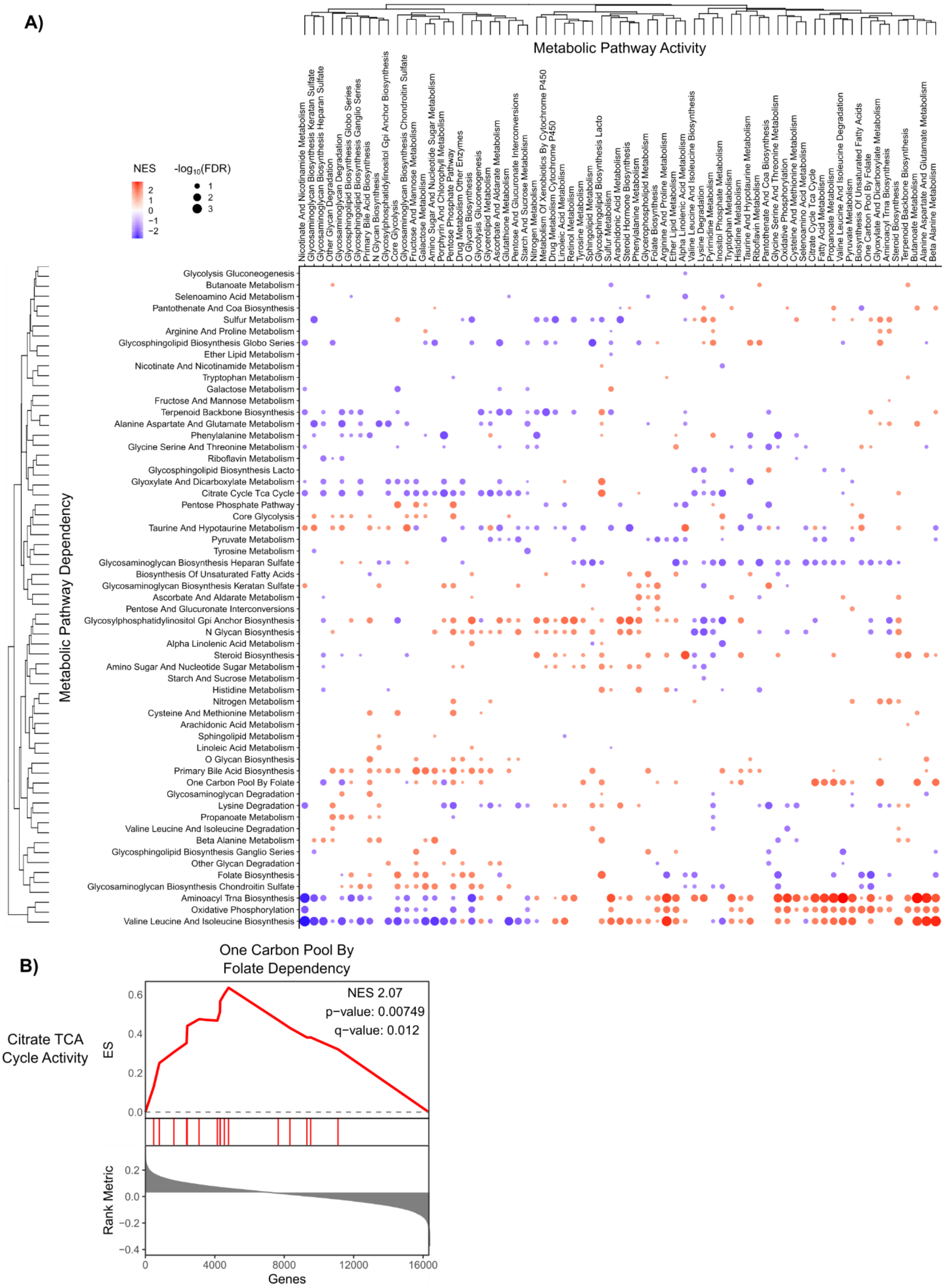
A)
Metabolic Pathway Activity
Nicotinate And Nicotinamide Metabolism
Glycosaminoglycan Biosynthesis Keratan Sulfate
Glycosaminoglycan Biosynthesis Heparan Sulfate
Other Glycan Degradation
Glycosaminoglycan Degradation
Glycosphingolipid Biosynthesis Globo Series
Glycosphingolipid Biosynthesis Ganglio Series
Primary Bile Acid Biosynthesis
N Glycan Biosynthesis
Glycosylphosphatidylinositol Gpi Anchor Biosynthesis
Core Glycolysis
Glycosaminoglycan Biosynthesis Chondroitin Sulfate
Fructose And Mannose Metabolism
Galactose Metabolism
Amino Sugar And Nucleotide Sugar Metabolism
Porphyrin And Chlorophyll Metabolism
Pentose Phosphate Pathway
Drug Metabolism Other Enzymes
O Glycan Biosynthesis
Glycolysis Gluconeogenesis
Glycerolipid Metabolism
Ascorbate And Aldarate Metabolism
Glutathione Metabolism
Pentose And Glucuronate Interconversions
Starch And Sucrose Metabolism
Nitrogen Metabolism
Metabolism Of Xenobiotics By Cytochrome P450
Drug Metabolism Cytochrome P450
Linoleic Acid Metabolism
Retinol Metabolism
Tyrosine Metabolism
Sphingolipid Metabolism
Glycosphingolipid Biosynthesis Lacto
Sulfur Metabolism
Arachidonic Acid Metabolism
Steroid Hormone Biosynthesis
Phenylalanine Metabolism
Glycerophospholipid Metabolism
Folate Biosynthesis
Arginine And Proline Metabolism
Ether Lipid Metabolism
Alpha Linolenic Acid Metabolism
Valine Leucine And Isoleucine Biosynthesis
Lysine Degradation
Pyrimidine Metabolism
Inositol Phosphate Metabolism
Tryptophan Metabolism
Histidine Metabolism
Taurine And Hypotaurine Metabolism
Riboflavin Metabolism
Pantothenate And Coa Biosynthesis
Glycine Serine And Threonine Metabolism
Oxidative Phosphorylation
Cysteine And Methionine Metabolism
Selenoamino Acid Metabolism
Citrate Cycle Tca Cycle
Fatty Acid Metabolism
Propanoate Metabolism
Valine Leucine And Isoleucine Degradation
Pyruvate Metabolism
Biosynthesis Of Unsaturated Fatty Acids
One Carbon Pool By Folate
Glyoxylate And Dicarboxylate Metabolism
Aminoacyl Trna Biosynthesis
Steroid Biosynthesis
Terpenoid Backbone Biosynthesis
Butanoate Metabolism
Alanine Aspartate And Glutamate Metabolism
Beta Alanine Metabolism
NES
2
1
0
−1
−2
-log10(FDR)
1
2
3
Metabolic Pathway Dependency
Glycolysis Gluconeogenesis
Butanoate Metabolism
Selenoamino Acid Metabolism
Pantothenate And Coa Biosynthesis
Sulfur Metabolism
Arginine And Proline Metabolism
Glycosphingolipid Biosynthesis Globo Series
Ether Lipid Metabolism
Nicotinate And Nicotinamide Metabolism
Tryptophan Metabolism
Galactose Metabolism
Fructose And Mannose Metabolism
Terpenoid Backbone Biosynthesis
Alanine Aspartate And Glutamate Metabolism
Phenylalanine Metabolism
Glycine Serine And Threonine Metabolism
Riboflavin Metabolism
Glycosphingolipid Biosynthesis Lacto
Glyoxylate And Dicarboxylate Metabolism
Citrate Cycle Tca Cycle
Pentose Phosphate Pathway
Core Glycolysis
Taurine And Hypotaurine Metabolism
Pyruvate Metabolism
Tyrosine Metabolism
Glycosaminoglycan Biosynthesis Heparan Sulfate
Biosynthesis Of Unsaturated Fatty Acids
Glycosaminoglycan Biosynthesis Keratan Sulfate
Ascorbate And Aldarate Metabolism
Pentose And Glucuronate Interconversions
Glycosylphosphatidylinositol Gpi Anchor Biosynthesis
N Glycan Biosynthesis
Alpha Linolenic Acid Metabolism
Steroid Biosynthesis
Amino Sugar And Nucleotide Sugar Metabolism
Starch And Sucrose Metabolism
Histidine Metabolism
Nitrogen Metabolism
Cysteine And Methionine Metabolism
Arachidonic Acid Metabolism
Sphingolipid Metabolism
Linoleic Acid Metabolism
O Glycan Biosynthesis
Primary Bile Acid Biosynthesis
One Carbon Pool By Folate
Glycosaminoglycan Degradation
Lysine Degradation
Propanoate Metabolism
Valine Leucine And Isoleucine Degradation
Beta Alanine Metabolism
Glycosphingolipid Biosynthesis Ganglio Series
Other Glycan Degradation
Folate Biosynthesis
Glycosaminoglycan Biosynthesis Chondroitin Sulfate
Aminoacyl Trna Biosynthesis
Oxidative Phosphorylation
Valine Leucine And Isoleucine Biosynthesis
B)
One Carbon Pool By
Folate Dependency
Citrate TCA
Cycle Activity
NES 2.07
p−value: 0.00749
q−value: 0.012
ES
0.6
0.4
0.2
0.0
Rank Metric
0.2
0.0
−0.2
−0.4
0
4000
8000
12000
16000
Genes

**Supporting Figure 2. Genetic PDEA in Adherent DMEM cell lines reveals context-specific pathway essentialities.** Metabolic pathway activity was inferred using single-sample GSEA (ssGSEA) for 153 adherent cell lines cultured in DMEM and correlated to gene dependency data from The Cancer Dependency Map (DepMap). Correlation coefficients were then ranked and GSEA was run querying the KEGG metabolic pathways (see Figure 1). **A)** Hierarchical clustering was performed on the Genetic PDEA normalized enrichment scores (NES). Results for pathways with FDR < 0.25 are plotted. Dots are colored according to their NES and sized according to the $-\log_{10}$ of the false discovery rate (FDR). Numerical values for each pathway can be found in Supp. Table 1. **B)** Increased TCA cycle activity is associated with increased dependency on One-Carbon Pool by Folate metabolism.

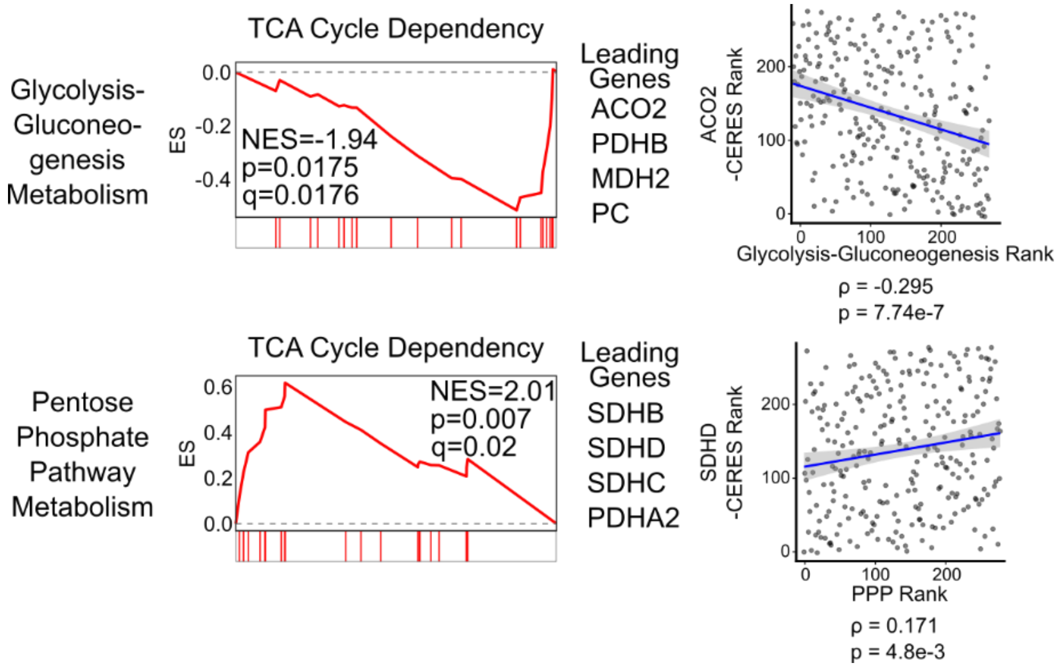


**Supporting Figure 3. Metabolic dependence on TCA cycle metabolism is context dependent.** In adherent RPMI cancer cells, when Glycolysis-Gluconeogenesis (hsa00010) pathway activity was low, the dependence on the TCA Cycle (hsa00020) was increased (top). When Pentose Phosphate Pathway activity (hsa00030) was high, the dependence on the TCA Cycle was increased (bottom). The scatter plots of pathway activity NES and gene dependency (-CERES) for leading-edge genes *ACO2* and *SDHD* with Glycolysis-Gluconeogenesis and Pentose Phosphate Pathway, respectively, are shown.

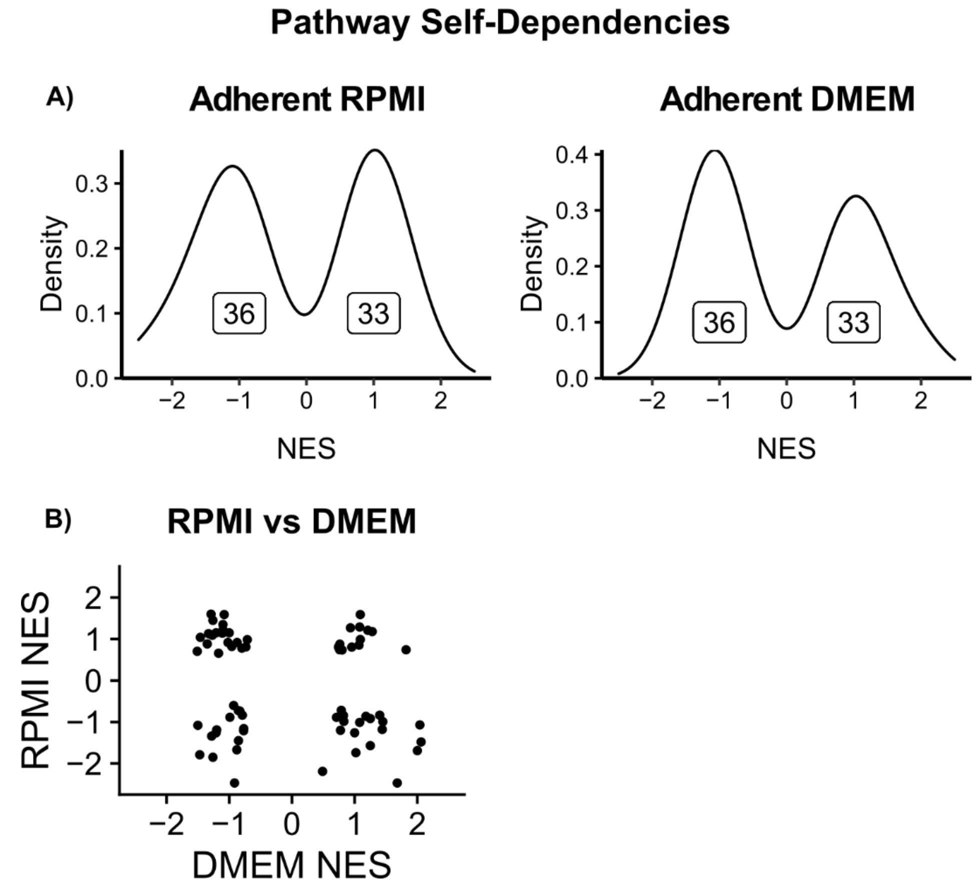


**Supporting Figure 4. Pathway activity does not correlate with pathway dependency.** Metabolic pathway expression was correlated with gene dependency data and GSEA was run on the resulting correlation coefficients (see Figure 1). Then, results were filtered for results that had the same pathway expression and pathway dependency tested (e.g. Glycolysis dependency GSEA was queried against all correlations with Glycolysis expression). **A)** The resulting normalized enrichment scores (NES) are presented as a density plot for both Adherent RPMI and Adherent DMEM analyses. The distribution of NES is centered around 0. **B)** The NES for each self-dependency is plotted for Adherent RPMI and Adherent DMEM. Values for each self-dependency can be found in Supp. Table 2.

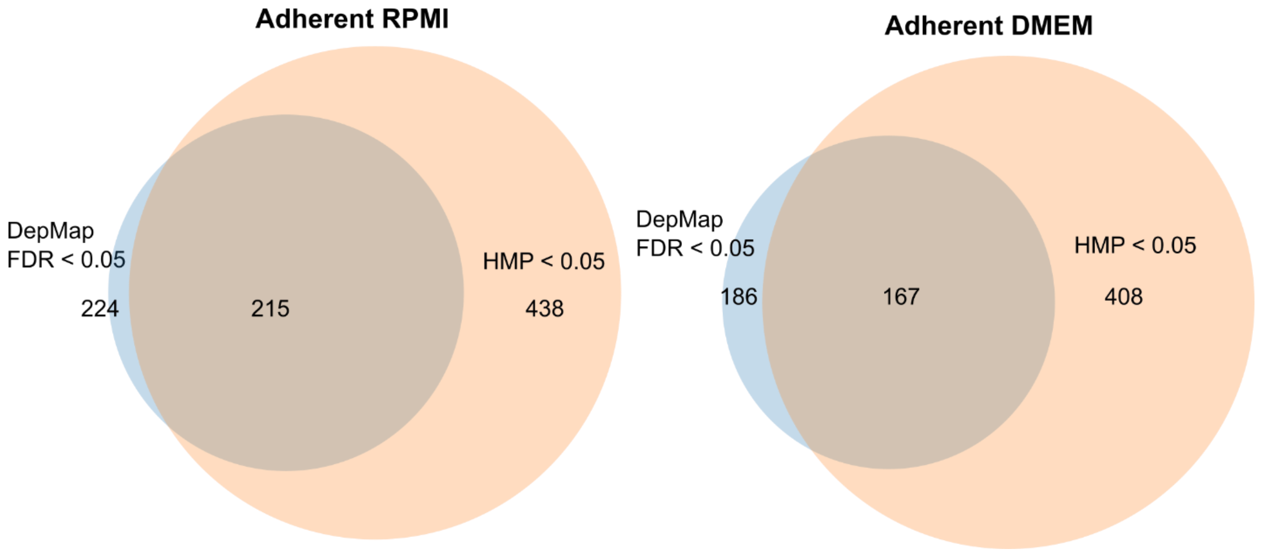


**Supporting Figure 5. Comparison of Genetic PDEA for gene dependency data sets between the Broad and Sanger Institutes.** To examine the reproducibility of Genetic PDEA, we compared data from pan-cancer CRISPR-Cas9 gene dependency data set (Sanger Institute) through the Genetic PDEA pipeline. We combined statistical tests using the harmonic mean p-value (HMP). When applying an HMP threshold of 0.05 and a false discovery rate threshold of 0.05, we see 96% and 90% agreement between the Genetic PDEA results from the DepMap and the Broad Institute.

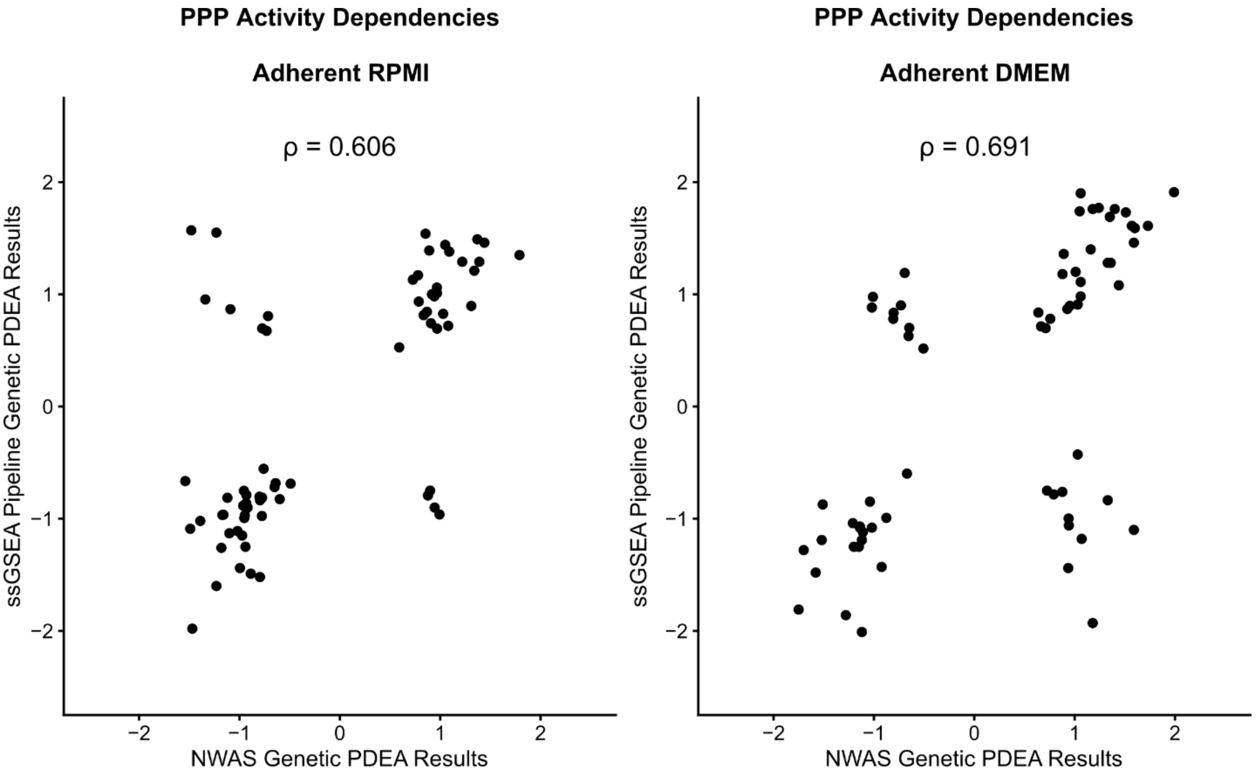


**Supporting Figure 6. Normalized weighted average expression (NWAS) gives similar results to ssGSEA for Genetic PDEA.** To directly compare NWAS with ssGSEA, we re-ran our pipeline using NWAS to analyze dependency on the Pentose Phosphate Pathway. We chose the Pentose Phosphate Pathway for this comparison because several enzymes are shared between pathways (e.g., PRKL, PFKM, and PKFP are present in both Glycolysis-Gluconeogenesis and Pentose Phosphate Pathway gene sets). We found broad agreement between the metabolic pathway dependencies when using either NWAS or ssGSEA for both Adherent RPMI and Adherent DMEM cell lines (Spearman r of 0.606 and 0.691, respectively).

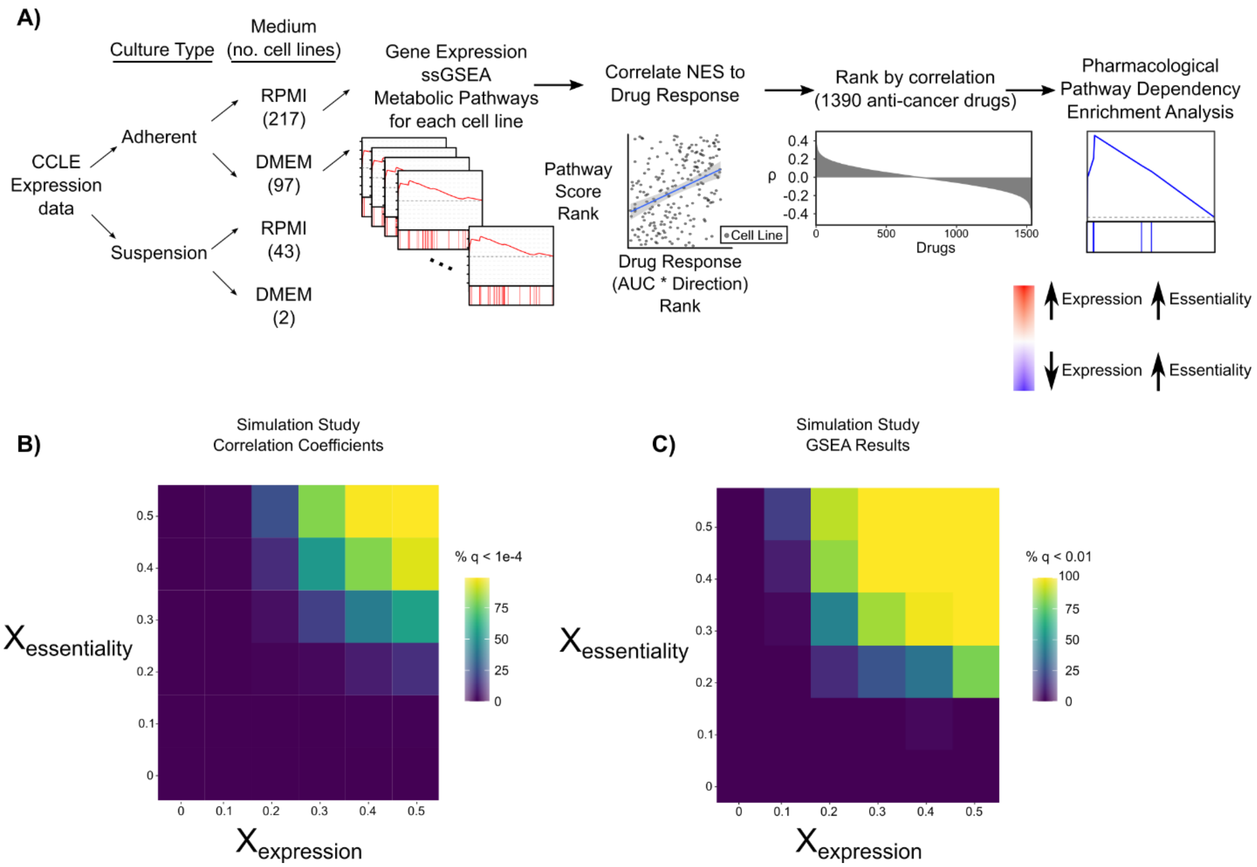


**Supporting Figure 7. Pharmacological Pathway Dependency Enrichment Analysis Simulation Study. A)** Schematic representing the strategy used to integrate metabolic pathway activity with drug response screens. Like Genetic PDEA (Fig. 1), the drug response (area-under-the-curve, AUC) for individual cancer cell lines was correlated to metabolic pathway activity as measured by ssGSEA. Drugs classified as activators (e.g., agonists) were multiplied by -1 for directional consistency. Cancer cell lines were separately processed by culture type and culture medium with a focus on adherent cell lines. All correlation p-values were FDR corrected using a Benjamini-Hochberg correction. Here, individual drug-pathway correlations are shown. Drugs were also mapped to metabolic pathways using their annotated gene targets and then Pharmacological PDEA was run on the resulting drug sets. Those results are presented in Fig. 5. **B-C)** Simulated data (see methods) was used to assess the sensitivity of the Pharmacological PDEA approach. Values added to the expression gradient resulted in slightly stronger correlation coefficients and Pharmacological PDEA results compared to values added to dependency gradient.

A)

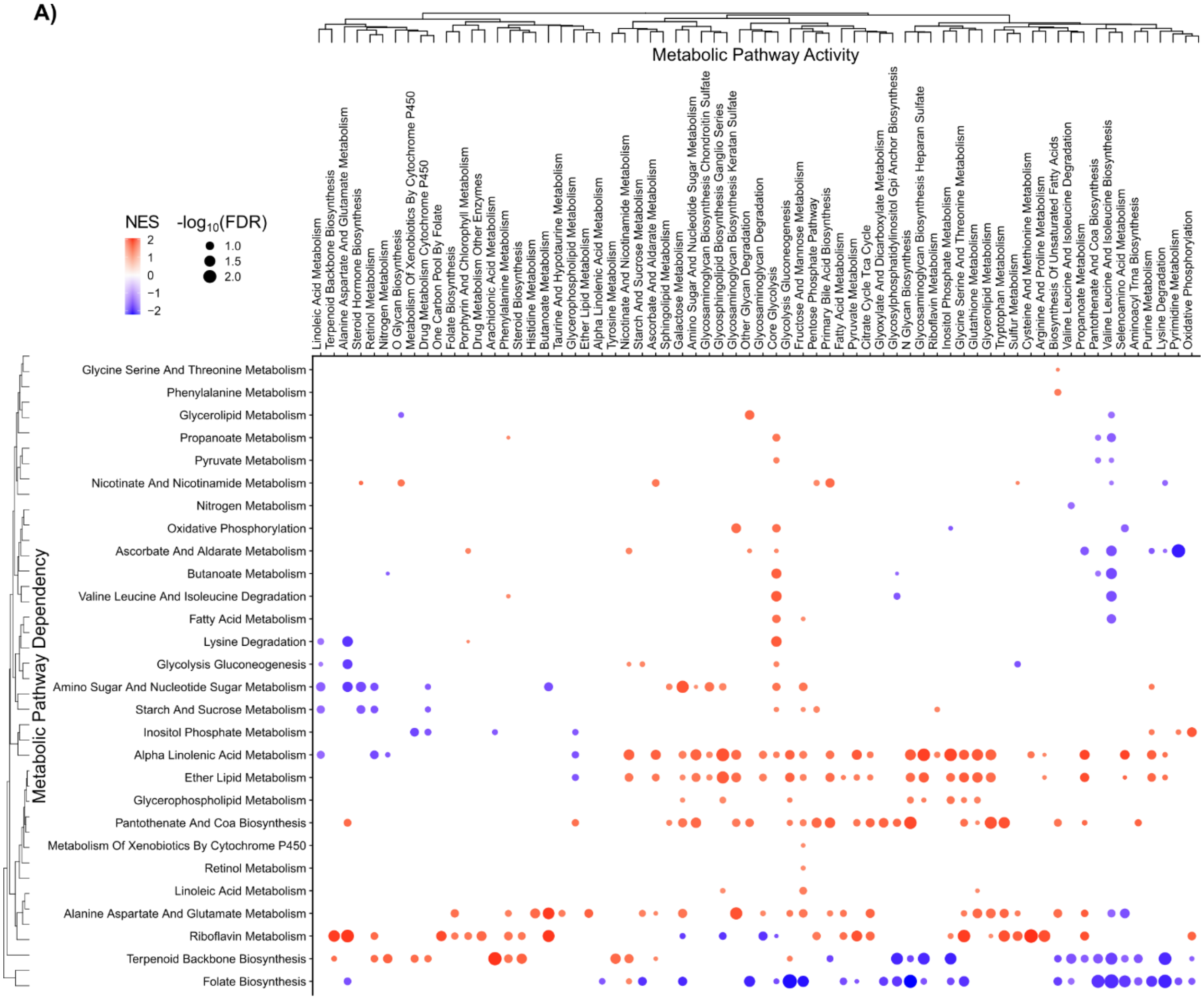


B)

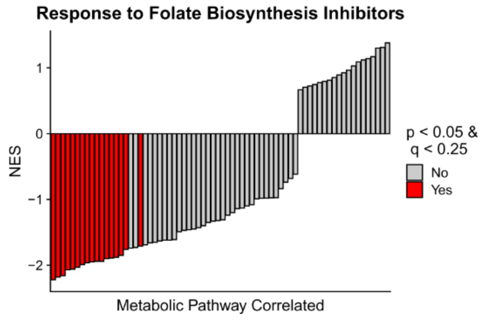


**Supporting Figure 8. Pharmacological PDEA Reveals Metabolic Pathway Vulnerabilities in Adherent DMEM cell lines**. **A)** Pharmacological PDEA (see Supp. Fig. 4) was performed on 1,390 anti-cancer drugs from the PRISM database for 97 adherent cell lines grown in DMEM. Drugs were mapped to metabolic pathways by their annotated target(s). Hierarchical clustering was performed on NES and results with FDR < 0.25 are plotted. Dots are colored according to the NES and sized according to the $-\log_{10}$ FDR. **B)** Inhibitors of Folate Biosynthesis (hsa00790) are more effective when overall metabolic pathway expression is low in Adherent DMEM cell lines.
